# The Consequences of Statistical Tests on Using Proxy Measurements in Place of Gold Standard Measurements: An Application to Magnetic Resonance Spectroscopy

**DOI:** 10.1101/2025.08.13.670023

**Authors:** Michael Treacy, Christoph Juchem, Karl Landheer

## Abstract

The use of proxy measurements in biomedical science is ubiquitous, due to the infeasibility or unavailability of gold-standard (i.e., most precise, accurate, and/or validated) measurements. For example, in magnetic resonance spectroscopy (MRS), short-echo time (TE) sequences are frequently employed to estimate difficult-to-measure metabolites such as GABA, despite J-difference editing being the recommended gold-standard for improved metabolic specificity. This work investigates the critical relationship between the correlation of proxy and gold-standard measurements and the associated false positive (FPR) and false negative (FNR) rates of statistical tests performed on proxy measurements. Through statistical simulations, we demonstrate that even moderately high correlations (0.6-0.7), reported in the literature for short-TE vs. J-edited estimated GABA, can lead to drastically inflated FPRs and FNRs. We show that these rates are highly sensitive to the magnitude of any introduced bias in the proxy measurement (δ) and the underlying true effect size (Δ). For instance, a small, unmeasured bias in short-TE estimated GABA, potentially arising from macromolecule contamination, can substantially inflate FPRs. Conversely, imperfect correlation can significantly reduce statistical power, leading to high FNRs, which may explain some discrepancies within the literature. Although this work focuses specifically on the relationship between short-TE and MEGA-edited GABA, the arguments presented here apply more broadly to other difficult-to-measure metabolites in MRS (e.g., glutathione, 2-hydroxyglutarate), or generally to *any* circumstance where statistical tests are performed on the readily available proxy measurements in place of gold-standard measurements.

## Introduction

In biomedical science, proxy measurements are widely used in place of gold-standard methods because the latter are often infeasible (due to resource or ethical constraints), impossible (due to inherent physical or technological limitations), or simply unavailable in existing datasets. In magnetic resonance spectroscopy (MRS), short-TE sequences are appealing for several factors: wide availability (preinstalled stock PRESS^1^ and STEAM^2^ sequences on all major vendor platforms), high SNR due to reduced T_2_- weighting^3^, reduced sensitivity to variations in T_2_ values, and easy quantification of typical major metabolites^3–5^ which may serve as a reference or may also be of interest. Short-TE sequences suffer from strong spectral overlap (Figure 1), and as such J-editing modules such as MEGA^6^ have been developed to improve metabolic specificty^7–9^, and have become the recommended standard^9,10^ for metabolites such as GABA, 2-hydroxyglutarate and glutathione. These metabolites are characterized by being low in amplitude and heavily overlapped with other signals, particularly macromolecules which have poorly characterized spectral shape^11^. Regardless, there are numerous examples of short-TE sequences being used to quantify GABA^12–22^, glutathione^13,15,17,23,24^ and 2-hydroxyglutarate^25–27^, to name a few.

**Figure 1.**
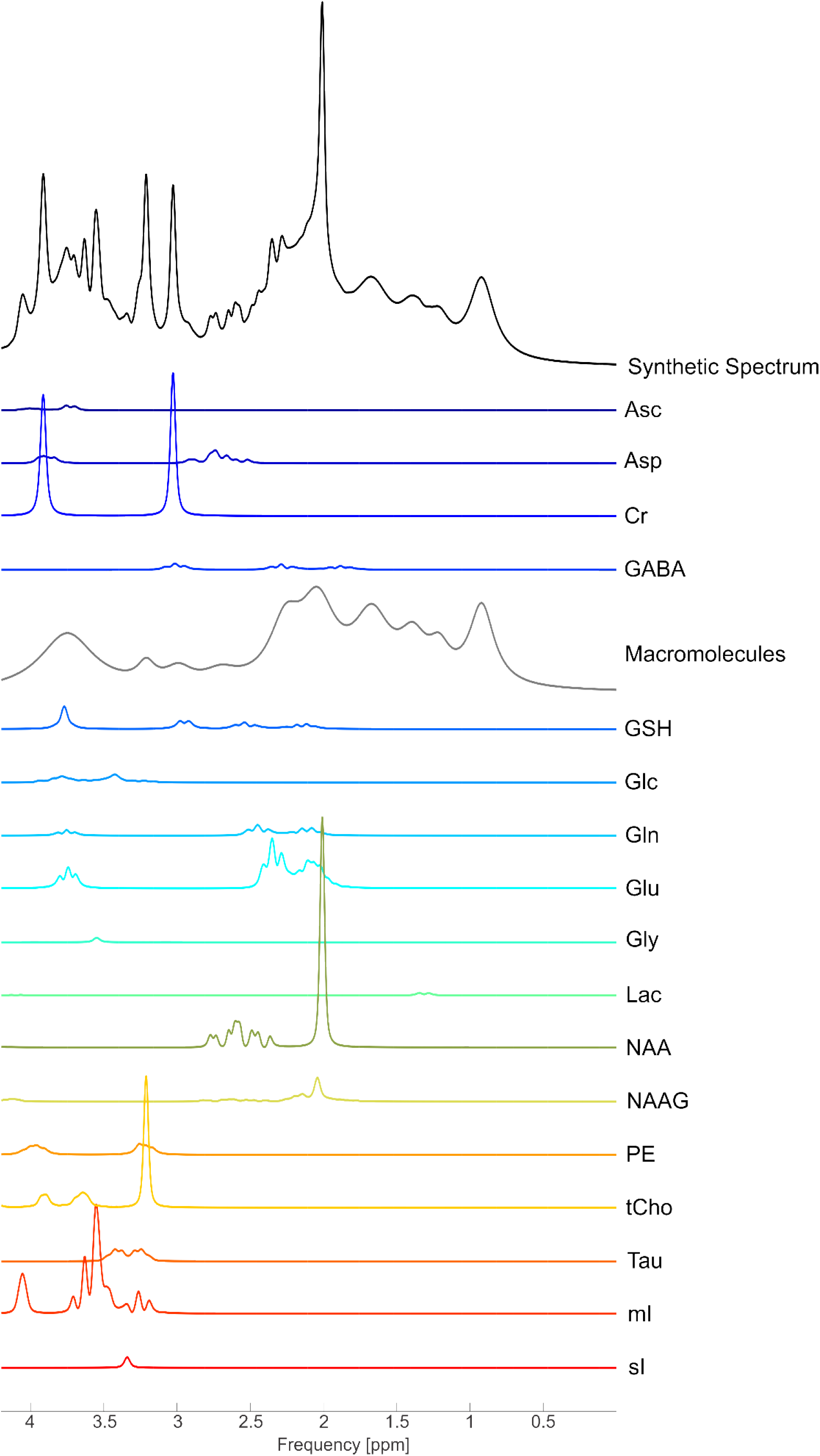
Synthetic spectrum of a typical short-TE (30 ms) PRESS sequence with repetition time = 2 s generated via synMARSS^45,46^ at 3T in the absence of noise for healthy brain tissue. Note the low relative amplitude of difficult-to-quantify metabolites such as GABA and GSH compared to numerous other components directly overlapping them, most notably the macromolecules. No individual moiety of either metabolite is free from spectral overlap with much more prominent resonances, which results in considerable quantification errors and moderate correlation with MEGA-edited quantification. Metabolites simulated include ascorbate (Asc), aspartate (Asp), creatine (Cr), γ-aminobutyric acid (GABA), macromolecules, glutathione (GSH), glucose (Glc), glutamine (Gln), glutamate (Glu), lactate (Lac), N-acetylaspartate (NAA), N-acetylaspartylglutamate (NAAG), phosphorylethanolamine (PE), total choline (tCho), taurine (Tau), myo-inositol (mI), scyllo-inositol (sI).

Provided that the correlation is sufficiently high between the gold-standard measurement (the focus of this work will assume this is MEGA-edited GABA) and the proxy measurement (i.e., short-TE measured GABA), simpler approaches may be appropriate. We investigate what exactly constitutes a “sufficiently high correlation” here. This is achieved using Monte Carlo statistical simulations, and we demonstrate that the false positive rate (FPR) and false negative rate (FNR) are strongly dependent on the correlation between measurements from MEGA- and short-TE estimate measurements. Even for the moderately-high correlations (0.6 to 0.7), representative of those from the literature for this application, the FPR and FNR can be drastically higher than reported, suggesting that results which rest on the assumption of perfect correlation need to be interpreted with increased caution^28,29^. A preliminary version of this work has previously been presented in abstract form^30^.

## Methods

We used Monte Carlo simulations to investigate the relationship between correlation of proxy variables to gold-standard variables and the FPR and FNR. We denoted A and B to be the random variables sampled from the gold-standard distributions (e.g., MEGA-edited measured concentrations for the metabolite of interest) of the control groups and cases, respectively. For example, B could represent concentrations from subjects in a disease population^31^, or concentrations after a functional stimulus^19^. Similarly, we denoted C and D as proxy variables (i.e., short-TE measured GABA concentrations) of the control and case population, respectively. The gold-standard measurements are modeled by

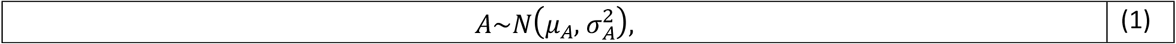

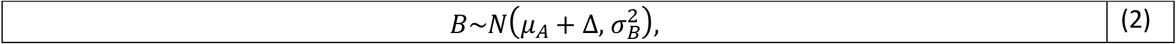

where 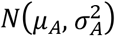 denotes the normal distribution with mean *μ*_*A*_ and standard deviation *σ*_*A*_, with unitsof millimol (mM) for concentrations, or unitless for relative concentrations, and ∼ indicates the sampling of such distributions. The true effect (i.e., differences in means between A and B) is given by Δ. We modeled the proxy measurements by

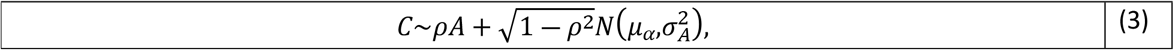

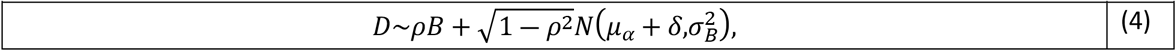

where 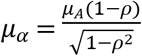, and a bias introduced by the use of proxy variables is given by *δ*. Derivations and assumptions for these expressions are given in the Appendix. Equations 1-4 exhibit the desired properties, namely: 1) *corr*(*A,C*)=*corr*(*B,D*)=*ρ*; 2) *var*(*A*)=*var*(*C*) and *var*(*B*)=*var*(*D*) (i.e., the use of proxy measurements does not change the variance); and 3) *E*(*C*)=*E*(*A*) (i.e., the proxy estimate is *unbiased* for the controls, but not necessarily the cases).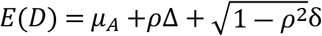, that is the mean of the proxy variable D contains a component of the true difference (*ρ*Δ), which has been dampened by the correlation coefficient, as well as a bias introduced that grows as correlation decreases 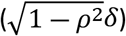.

We calculated the false positive and false negative rates numerically in accordance with the above equations. All simulations were performed with 500,000 times per set of parameters with NumPy version 1.21.6 and all statistical tests were two-sided independent Welch’s t-tests performed via SciPy version 1.7.3. We performed statistical tests on the proxy variables (i.e., C and D), under the assumption that we are actually interested in detecting whether Δ ≠ 0 (i.e., a difference in the gold-standard variables). The code to reproduce all statistical simulations is available at: github.com/karllandheer/GoldStandardVsProxy.

To investigate FPRs, we set Δ= 0 (i.e., null hypothesis is true, no difference between the means of the gold-standard distributions) and *δ* > 0, i.e., a bias is introduced in the estimation of cases but not controls of the proxy variables; possible scenarios where this could arise are explored in the discussion. In short, this scenario arises when one is measuring a difference between cases and controls that is assumed to be due to a change in GABA concentration, but is actually due to a different reason. To investigate FNRs, we set *δ*= 0 (no bias) but Δ > 0 (null hypothesis is false). Initially, we array across a large range of possibilities common in biomedical literature to demonstrate that in certain scenarios both FPR and FNR can be strongly sensitive to correlation. These parameters include nominal false-positive rates *α* ∈ {0.05, 0.01, and 0.001}, number of samples, *N*_*samples*_ ∈ {10, 25, 100}, all possible positive correlations (from 0 to 1). While *δ* and Δ values are unmeasured, we include a range large enough to demonstrate the relationship between FNR and FPR with correlation.

We then performed statistical simulations with experimental literature values. First, we used values from the literature which report statistically significant difference in the concentration of GABA in the visual cortex before and after behavioral training as assessed by short-TE MRS at 7T^19^. We assume the sample values are reflective of the population values, which includes, i.e., *σ*_*A*_= 1.55, *σ*_*B*_=2. 05, *α*= 0. 05, *μ*_*C*_=6.8, *μ*_*D*_=7.6. This study had a number of samples N_samples_ = 25 and a nominal false positive rate of *α*= 0. 05. We calculated what *δ* may explain the observed differences between distributions and provide the mathematical context of how such a *δ* may arise due to small macromolecule contamination.

To investigate the influence of correlation on FNR, we then used values from the literature which used short-TE to estimate GABA at 3T but failed to find a difference between ALS subjects^12^, and investigate what Δ would be required to produce a well-powered study (i.e., FNR = 0.05). We showed that such a value is well below the value previously found in an experiment which detected a true difference in GABA in ALS and controls with a MEGA-edited sequence^32^. The relevant values from this study are *N*_*samples*_=9/1 0 (cases/controls), and once again assume the population means and standard deviations are equal to the sample values, *μ*_*C*_= 0.312, *σ*_*c*_= 0. 06, *σ*_*D*_= 0. 02, and *α*= 0. 05. Previously, it was observed that ALS patients had a 16% reduction in GABA (1.42 in ALS patients vs 1.70 in healthy controls^32^), thus we assumed the true effect size is Δ=− 0. 05.

## Results

In the case when Δ= 0, FPR increases rapidly with correlations that deviate from unity if the proxy-measurement bias, *δ*, is large (Figure 2). At even moderate to strong correlations (0.4 to 0.7), provided *δ* is sufficiently large, the FPR can be above 80%. The false positive rate *increases* with increasing sample size, as the experiment is better powered to detect a false result. Importantly, the FPR decreases with decreasing *α*, as it requires a more stringent cutoff to detect a statistically significant result. This is one potential strategy to mitigate such an effect, although it invariably comes with the decreased ability to detect true associations as well.

**Figure 2.**
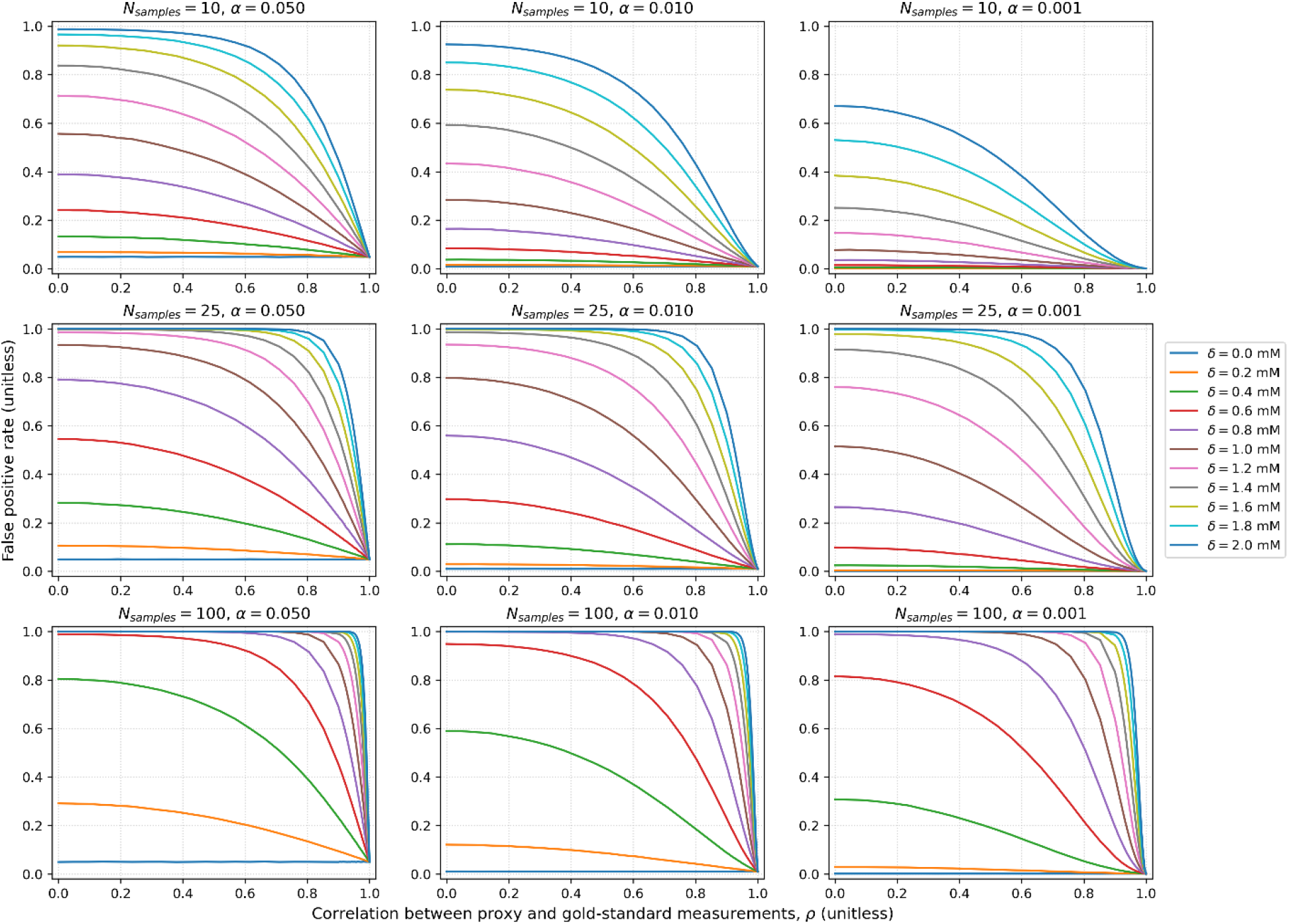
False positive rate vs correlation across a range of bias values (*δ*). Values were chosen of *σ*_*A*_= *σ*_*B*_=1 *mM* and *μ*_*A*_= 2.1 mM (typical concentration for GABA^3,39^). An array across a typical number of subjects within the neuroimaging literature, N_samples_ = 10 (top row), N_samples_ = 25 (middle row) and N_samples_ = 100 (bottom row). Nominal false positive error rates are displayed with *α*= 0. 05 (left column), *α*= 0. 01 (middle column) and *α*= 0. 0 01. Even moderate to strong correlations (0.4 to 0.7), provided *δ* is sufficiently large, the false-positive rate can be nearly 100%. The false positive rate *increases* with increasing sample size, as the experiment is better powered to detect a false result (i.e., the proxy-induced bias). Important to note, the false positive rate decreases with decreasing *α*, as it requires a more stringent cutoff in order to detect a statistically significant result. At a correlation of 1.0 the false positive rate converges to the nominal *α*, as expected.

In the case when Δ ≠ 0, but *δ*= 0, even with moderate to strong correlation between gold-standard and proxy measurements, the use of proxy measurements results in a substantial increase in FNR (Figure 3). For example, with *N*_*samples*_ = 25, *α*= 0. 01 (center plot), if Δ=1.6 *mM*, the false negative rate can be approximately 10%, while at a correlation of 0.7 it’s nearly 40%. The FNR increases with increasing *α*, as expected, and decreases with *N*_*samples*_.

**Figure 3.**
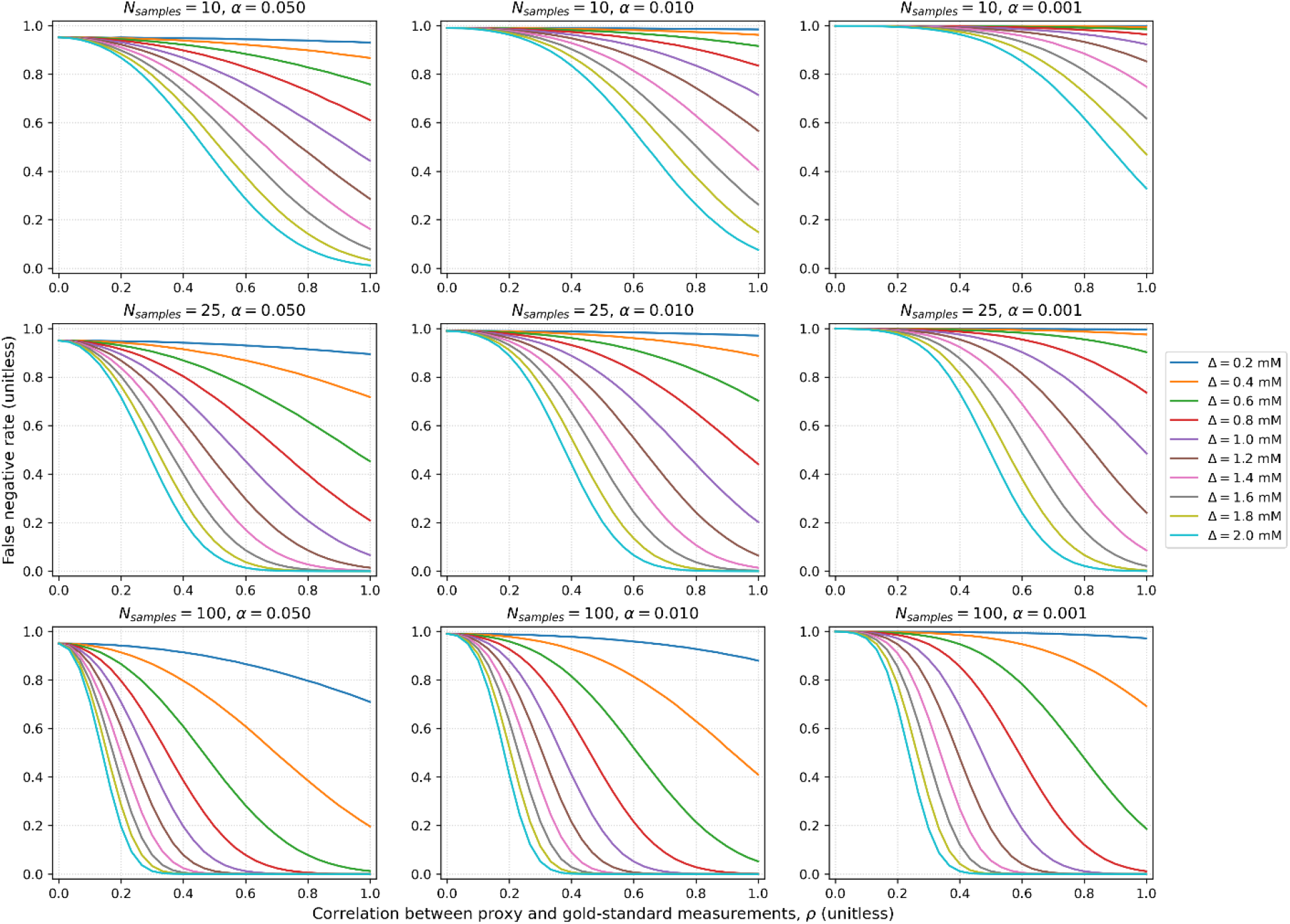
False negative rate vs correlation across a range of differences between gold standard measurements (Δ). Values were chosen of *σ*_*A*_= *σ*_*B*_=1 *mM* and *μ*_*A*_= 2.1 mM (typical concentration for GABA^3,39^). An array across a typical number of subjects within the neuroimaging literature, N_samples_ = 10 (top row), N_samples_ = 25 (middle row) and N_samples_ = 100 (bottom row). Nominal false positive error rates are displayed with *α*= 0. 05 (left column), *α*= 0. 01 (middle column) and *α*= 0. 0 01. Even with moderate-to-strong correlations the false negative rate can be substantially inflated. For example, with N_samples_ = 25, *α*= 0. 01 (center plot), if Δ=1.6 *mM* at a correlation of 1 the false negative rate is approximately 10%, while at a correlation of 0.7 it’s nearly 40%. The false negative rate increases with increasing *α* and decreases with *N*_*samples*_, as expected. At low correlations regardless of differences between the true distributions the false negative rate converges to the value of 1 − *α*, as expected.

Using experimental values from the literature^19^, we demonstrate that the FPR increases dramatically with *δ*. While such *δ* values are unmeasured, we provide an intuition in the Discussion that the values required to substantially inflate the FPR could easily be the result of small variations in macromolecule concentrations between cases and controls. If we assume the null hypothesis is correct (i.e., Δ= 0), and that the means of the measured values are their true parameters, then 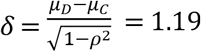. With *δ*=1.19 the FPR is 35%, substantially higher than their nominal chosen value of 5%, indicating this result should be interpreted with increased caution.

Using values from the literature^12^, we show that the use of proxy variables can substantially reduce statistical power. Using their measured values and a correlation of 0.58 (as previously measured at 3T^16^), the required Δ to achieve a FNR of 5% is −0.14. The true effect size was assumed to be a 16% reduction in GABA/creatine (hence Δ=− 0. 05), as previously found in a MEGA-edited experiment^32^ in the same disease. This 16% reduction results in a Δ=− 0. 05, roughly three times smaller than what is required to achieve a FNR of 0.05. With Δ=− 0. 05, the false negative rate is 72%. Meanwhile, if the correlation was 1 (i.e., the use of MEGA-editing to measure GABA), the required effect size to achieve a FNR of 5% is reduced by nearly a factor of two to Δ=− 0. 08. With a correlation of 1 and their measured value of Δ=− 0. 05 the FNR is 33%. Thus, this apparent discrepancy between Bilcher, et al.^12^ and Foerster, et al.^32^ may be explained by the study by Bilcher et al. being underpowered, in part due to use of short-TE to measure GABA which reduces statistical power, which can also be immediately observed from Figure 3.

## Discussion

This work suggests a comprehensive characterization of the relationship between proxy and gold-standard methods is critical for statistical interpretation of the experiments using the former. We demonstrated via Monte Carlo simulations that typical correlations found in the literature between short-TE and MEGA-edited GABA do not preclude the possibility of drastically increased false positive and false negative rates. We emphasize that, while this work has made numerous assumptions (as detailed in the Appendix), such assumptions result in the desired correlations between proxy and gold-standard measurements, which is partially the basis for the validity of such short-TE measurement results. We have shown across a wide range of parameters that the use of imperfectly correlated proxy measurements can result in the substantial inflation of false positive and false negative rates. Whether this argument impacts specific experiments depends on parameters not currently measured (i.e., *δ* and Δ), thus the focus of this work is simply to bring attention to this issue.

It is important to note that while this work focused on the MEGA-edited experiments as the purported gold standard for certain metabolites^9^, MEGA-edited measurements are themselves a proxy for true concentrations. Thus, this work also has implications for the interpretation of MEGA-edited measurements. Currently, high-quality independent validation measurements of MEGA-edited MRS with other MR or non-MR methods are limited. Selected ex vivo validation of post-mortem tissues have been performed preclinically^33^ and clinically^34^. For example, in one animal study, the correlation of short-TE measured GABA with ex vivo concentrations from enzyme-linked immunosorbent assay was 0.66 at 7T^33^, once again in the range where FPRs and FNRs can be inflated. Statistical advancements are being developed to address the related issue of how to best combine available proxy and gold-standard measurements^35^. Typical quantified errors do not address the issue investigated here for two important reasons: firstly, and most importantly, we demonstrate that one can be measuring effects (e.g., macromolecules) entirely distinct from the effects one believes they are measuring (GABA), even while maintaining medium to high correlations with the desired quantity. Secondly, estimation of uncertainty is invariably difficult in MRS, as typical estimates via the Cramér-Rao Lower Bound require knowledge of the underlying model to fit the experimental data to, which is not known^36^, and distribution-free methods such as Conformal Predictors require access to underlying ground truth examples^37,38^, which are also inherently unavailable.

To estimate the increase in FPR over the nominal value the proxy-induced bias, *δ*, must be known. While such values are experimentally unmeasured, the calculation below shows that a small variation in macromolecule concentration, due to their substantially higher amplitudes at short-TEs, could result in bias values comparable to those used in simulations in this work. While GABA has a concentration of approximately 2.1 mM in the healthy brain^3,39^, the overlying macromolecules have an absolute proton concentration, *C*_*M,abs*_, of 19.8 mM at 2.99 ppm, 120.8 mM at 2.04 ppm, and 49.0 mM at 2.26 ppm, which nearly directly overlap with the GABA moieties at 3.01 ppm, 1.89 ppm and 2.23 ppm, respectively. Thus, using the institutional units where GABA/tCr ≅ 7.6 from Jia *et al*.^19^ (referred to as *C*_*GABA,inst*_) and absolute concentration of GABA is 2.1 *mM* (referred to as *C*_*GABA,abs*_), the institutional concentrations of these three macromolecule resonances would be 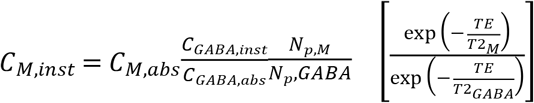, where the 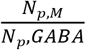 corrects for *C*_*M,abs*_ being a proton concentration^11^, and *C*_*GABA,abs*_ being a molecular concentration (this factor is equal to ½ for all resonances as all moieties of GABA contain 2 protons^40^) and the last term corrects for the difference in T_2_ values between GABA and macromolecules at the experimental TE of 36 ms. T_2_ values of 63 ms were used for GABA (measured at 7T^41^), and 20.0 ms, 14.3 ms, 19.8 ms, for M_2.99_, M_2.04_, M_2.26_, (measured at 3T^11^ with similar values obtained at 9.4T^42^). Thus, 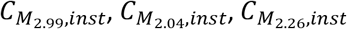 are 10.5, 31,2, and 25.5 (unitless), respectively. Thus, if even a small change (i.e., a few %) in these macromolecules concentrations between conditions bled into the GABA signal, it would result in a *δ* which could explain the observed difference of 1.19 (Figure 4). While Jia *et al*.^19^ used an inversion preparation module to measure the macromolecules, this only partially alleviates the issue as the macromolecule measurement can vary substantially from those in the short-TE spectrum. This is because the inversion preparation module heavily weights the T_1_ spectrum^11^. Furthermore, although macromolecules are most problematic due to their poorly characterized spectral shape, there are numerous other directly overlapped metabolites with GABA, such as NAA, NAAG, GSH, glutamate (Glu) and glutamine (Gln) (Figure 1) which could also potentially bleed into the estimated GABA quantity. While macromolecules are a likely contender for the introduction of such bias, other experimental differences between cases and controls could also cause it. Potential causes include differences in shim quality due to tissue heterogeneity differences^43^ (e.g., a tumor) or motion differences (e.g., Alzheimer’s^44^).

**Figure 4.**
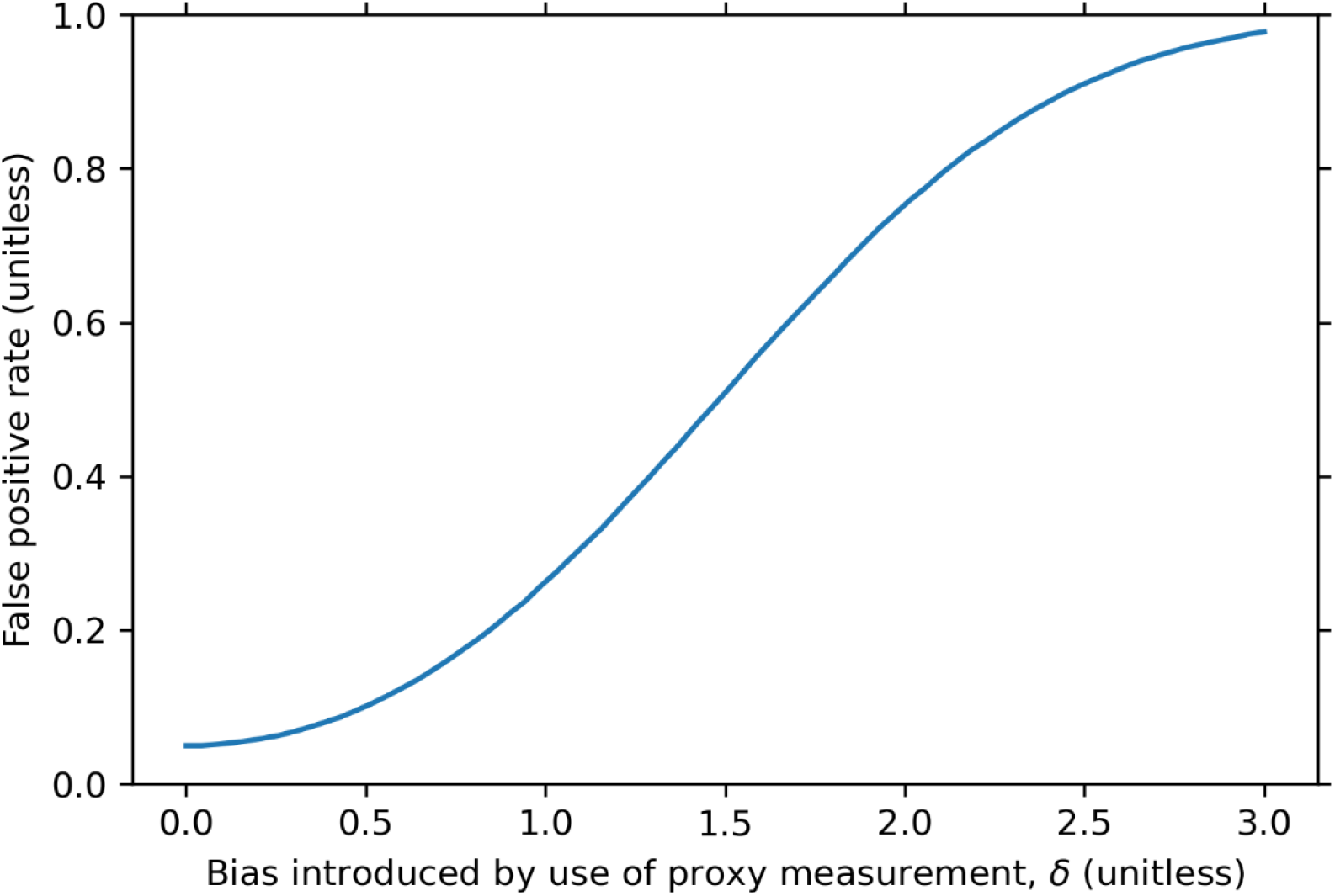
False positive rate versus *δ* with experimental values claiming to find a statistically significant difference in GABA/Creatine before and after behavioral training^19^ (n = 25 for both cases/controls). Reported values^19^ of *σ*_*C*_=1.55, *σ*_*D*_=2. 05 were used. Assuming the null hypothesis is true, the difference in distributions could be explained by, δ=1.19 (calculated using the empirical correlation of 0.72 at 7T^47^, and the reported difference between proxy measurement of *μ*_*D*_ − *μ*_*C*_= 0.83). Using this *δ*, at a correlation of 0.72 the FPR was calculated to be 35%, substantially higher than the nominal chosen value of 5%, thus their purported result should be interpreted with increased caution.

**Figure 5.**
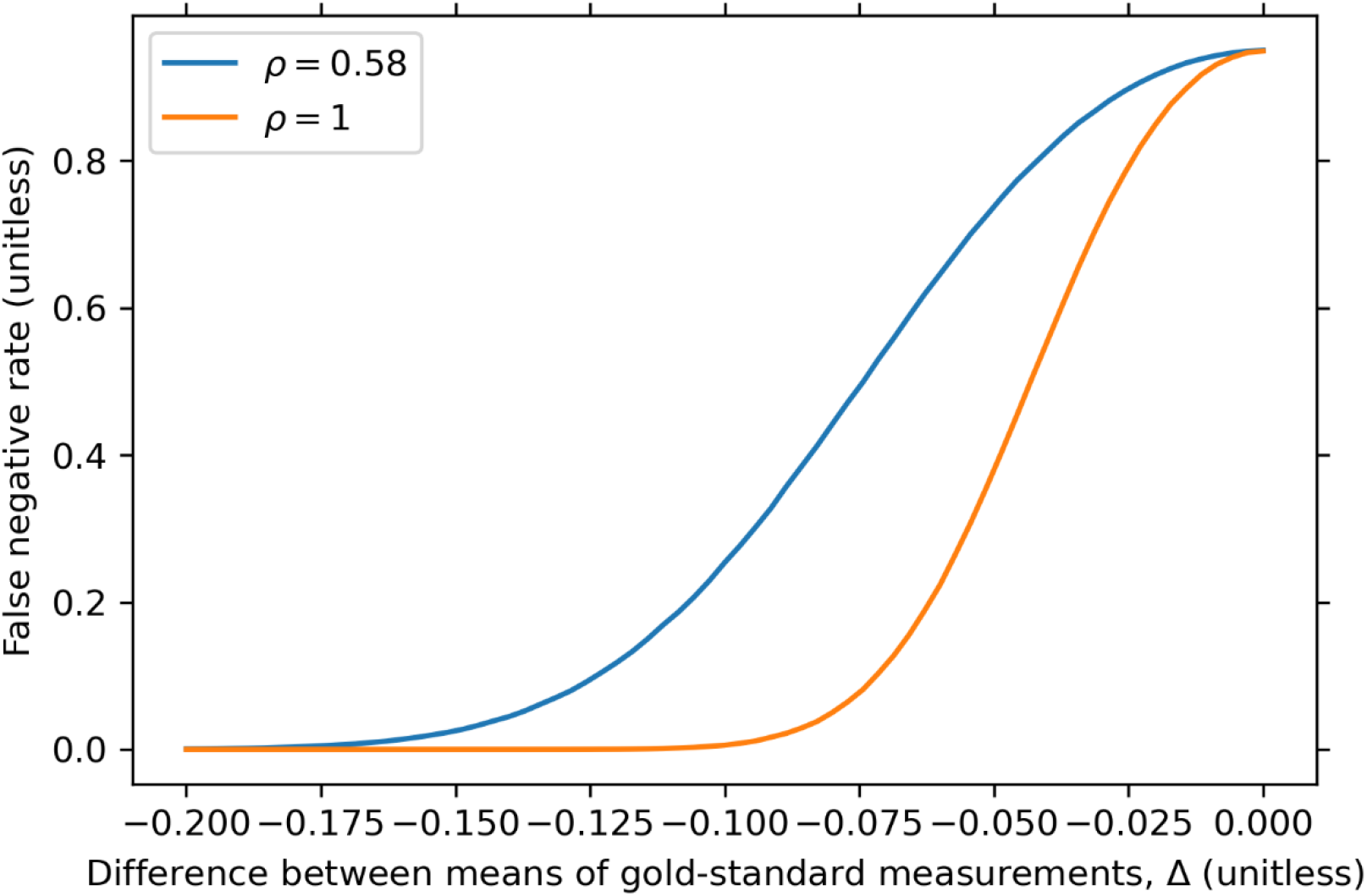
False negative rate versus Δ using experimental values unable to find a previously reported difference in GABA/creatine between ALS patients (n = 9) and healthy controls^12^ (n = 10), with *μ*_*A*_= 0.312, *σ*_*C*_= 0. 06 and *σ*_*D*_= 0. 02. The true effect size was assumed to be a 16% reduction in GABA/creatine (hence Δ=− 0. 05), as previously found in a MEGA-edited experiment^32^. For a correlation of 0.58 at 3T^16^ the false negative rate was calculated to be 72%, meanwhile with a correlation of 1.0 (i.e., gold standard), the false negative rate would be 35%, indicating that while the use of proxy measurements drastically reduces the power of detecting a difference for this study, the null result could also be easily explained by simply being underpowered even with the use of gold-standard measurements.

One limitation of the work presented here is the assumption of normality of distributions of the random variables. While this is a common assumption used within the biomedical literature (hence the use of t-tests), there is no inherent physiological reason why such concentrations would be sampled from a normal distribution. We chose the normal distribution as, for a given mean and standard deviation, it maximizes the entropy of the random variables. Hence in the absence of additional information about the distributions it becomes a natural choice. Work on characterizing the distributions of metabolite concentrations from both healthy and diseased populations would be a valuable contribution and could be used to further refine the results obtained here, among other simulations. Regardless, while the exact numerical results will invariably change based on choice of distributions, the problem investigated here will persist.

## Acknowledgements

Special thanks to Dr. Kelley Swanberg for fruitful discussions regarding the material of this work, and Dr. Helge Jörn Zöllner for encouraging its publication. This work was conducted independently and is unrelated to the Karl Landheer’s employment at Regeneron Pharmaceuticals, Inc.

## Appendix

In this appendix, we derive the expressions that are used in the statistical simulations. To obtain equations 3 and 4, the goal is to define the proxy variable as a linear combination of the gold-standard variable, *Y*_1_, and an independent noise term, *Y*_2_, that exhibit the desired correlation, i.e., *Y*_1_=*x*_1_*Y*_1_ +*x*_2_*Y*_2_. The following assumptions were made: 1) The gold-standard variable, *Y*_1_, and the *noise* term from the proxy measurement, *Y*_2_, are independent, i.e., *cov*(*Y*_1_,*Y*_2_)= 0. 2) the gold-standard variable and the noise term have equal variance, i.e., *var*(*Y*_1_)=*var*(*Y*_2_). 3) the variance of the proxy variable is equal to the variance of the gold-standard variable, i.e., *var*(*Y*_1_)=*var*(*x*_1_*Y*_1_ + *x*_2_*Y*_2_). First, we demonstrate how equations 3 and 4 arise from these assumptions.

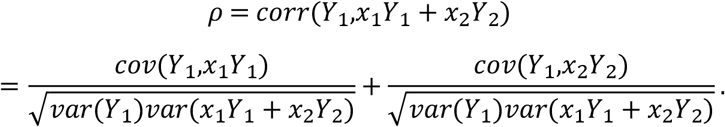

By assumption 1) the right-hand term becomes 0, thus and subbing in assumption 2 this becomes

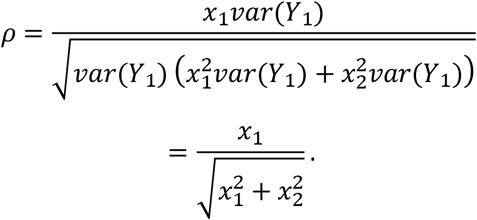

Assumption 3 states that 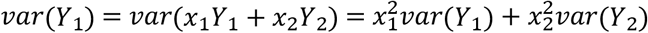. Thus, by assumption 2) this becomes 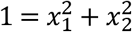. Subbing this into the expression above, we obtain *x*_1_=*ρ*,which can be used to derive 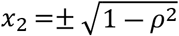. This can be used to immediately derive equations 3 and 4, where we have chosen the positive solution, 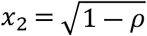, (the choice of positive or negative is arbitrary as we employed two-sided tests).

Below we show that *E*(*C*)=*E*(*A*) (i.e., proxy variables are unbiased estimators of gold standard variables):

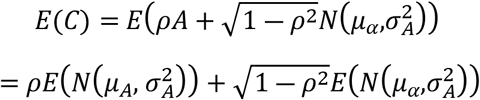

We set 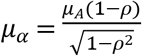,

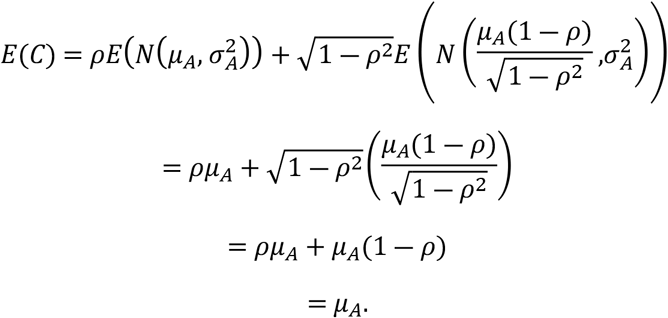

Finally, we show that 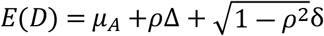. Starting with

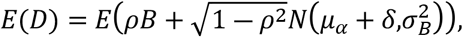

we sub in the definitions of *B* and 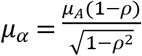 to obtain

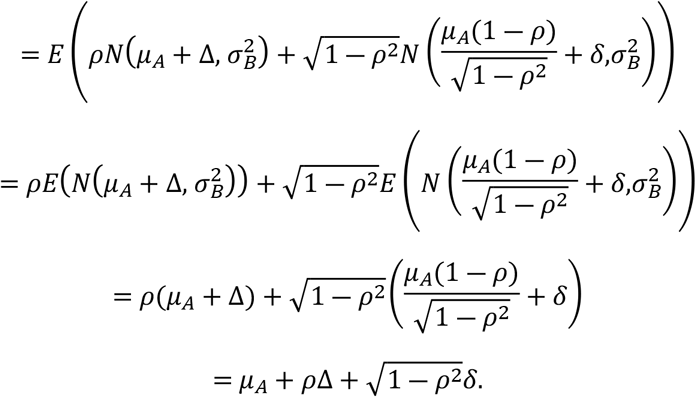

